# T cells limit accumulation of aggregate pathology following intrastriatal injection of α-synuclein fibrils

**DOI:** 10.1101/2020.02.20.956599

**Authors:** Sonia George, Trevor Tyson, Nolwen L. Rey, Rachael Sheridan, Wouter Peelaerts, Katelyn Becker, Emily Schulz, Lindsay Meyerdirk, Amanda R. Burmeister, Jennifer A. Steiner, Martha L. Escobar Galvis, Jiyan Ma, Andrew Pospisilik, Viviane Labrie, Lena Brundin, Patrik Brundin

## Abstract

Background: α-Synuclein (α-syn) is the predominant protein in Lewy-body inclusions, which are pathological hallmarks of α-synucleinopathies, such as Parkinson’s disease (PD) and multiple system atrophy (MSA). Other hallmarks include activation of microglia, elevation of pro-inflammatory cytokines, as well as the activation of T and B cells. These immune changes point towards a dysregulation of both the innate and the adaptive immune system. T cells have been shown to recognize epitopes derived from α-syn and altered populations of T cells have been found in PD and MSA patients, providing evidence that these cells can be key to the pathogenesis of the disease. Objective: To study the role of the adaptive immune system with respect to α-syn pathology. Methods: We injected human α-syn preformed fibrils (PFFs) into the striatum of immunocompromised mice (NSG) and assessed accumulation of phosphorylated α-syn pathology, proteinase K-resistant α-syn pathology and microgliosis in the striatum, substantia nigra and frontal cortex. We also assessed the impact of adoptive transfer of naïve T and B cells into PFF-injected immunocompromised mice. Results: Compared to wildtype mice, NSG mice had an 8-fold increase in phosphorylated α-syn pathology in the substantia nigra. Reconstituting the T cell population decreased the accumulation of phosphorylated α-syn pathology and resulted in persistent microgliosis in the striatum when compared to non-transplanted mice. Conclusion: Our work provides evidence that T cells play a role in the pathogenesis of experimental α-synucleinopathy.

## INTRODUCTION

The pathogenesis of Parkinson’s disease (PD) and multiple system atrophy (MSA) are poorly understood, but neuroinflammation and α-synuclein (α-syn) aggregation are believed to play key roles[1–4]. Progressive spread of Lewy pathology is thought to contribute to clinical decline in PD and to be the result of cell-to-cell propagation of α-syn aggregates[5]. Neuroinflammation is believed to be involved in the initial formation and spread of α-syn aggregates[4,6–8]. Studies indicate that activated microglia and elevated neuroinflammatory markers in the central nervous system are present in MSA and PD[3,9–15]. Single nucleotide polymorphisms close to numerous immune system-related genes affect PD risk, further supporting a role for neuroinflammation in PD[16]. While the majority of studies exploring inflammation in PD have implicated changes in the innate immune system[4,17], the role of the adaptive immune system in PD has been explored to a lesser extent. Notably, peripheral immune cells enter the brain during neurodegeneration[18]. T lymphocytes are altered and infiltrate the brain in PD[19–26] and it has been reported that autoreactive T lymphocytes directed against α-syn are present in PD patients[27] decades prior to motor PD[28]. Recruitment of CD4+ T cells to the brain occurs in models of a-syn overexpression[29]. In PD models, T cell function has been linked to α-syn pathobiology[30–32] and to the death of dopamine neurons[8,33]. Specifically, T cells respond to α-syn variants associated with PD and then migrate into the brain where they affect the phenotype of microglia[32]. Overexpression of α-syn induces microglia to express major histocompatibility complex II (MHC II)[34] and to present antigens to CD4^+^ T cells[8,33]. These interactions between T cells, microglia, and pathogenic α-syn are, however, not well understood. It has been difficult to interrogate the roles of the different populations of immune cells in a model of α-synucleinopathy, as it has been done for other diseases (e.g., Multiple Sclerosis[35]). We addressed this gap in knowledge by triggering α-syn pathology in immunocompromised (NOD scid gamma: NSG, lacking T, B and Natural Killer cells) mice and reconstituting select populations of immune cells. Specifically, we triggered α-syn pathology by intrastriatal injection of human α-syn preformed fibrils (PFFs) in control mice, NSG mice and NSG mice where T or B cell populations had been reconstituted. We assessed pathological accumulation of phosphorylated α-syn in multiple brain regions and found that NSG mice displayed elevated pathology, while the reconstitution of T cells in NSG mice was associated with partial reduction of α-syn neuropathology.

## MATERIALS AND METHODS

### Study design

The goal of our study was to define whether a compromised immune system influences the accumulation of pathological α-syn in the brain *in vivo*. To this end, we assessed phosphorylated α-syn pathology following intrastriatal injections of human α-syn PFFs in immunocompromised and control mice. First, we established whether the PFFs injection in the striatum of immunocompromised mice resulted in altered α-syn pathology compared to when the PFFs were injected into wildtype mice. Second, we determined the effects on neuropathology of reconstituting T cells in immunocompromised mice that had received intrastriatal PFFs.

### Animals

We utilized 10 to 12-week-old female C57BL/6J, NOD/ShiLtJ (non-obese diabetic) and NSG mice [immunocompromised, lacking T, B and Natural Killer (NK) cells] mice for injections, and additionally isolated T and B cells from female C57BL/6J mice (bred in our internal vivarium colony). Mice were housed at a maximum of five per cage under a 12-h light/12-h dark cycle with access to food and water *ad libitum*. The housing of the animals and all procedures were carried out in accordance with the *Guide for the Care and Use of Laboratory Animals* (United States National Institutes of Health) and were approved by the Van Andel Research Institute’s Institutional Animal Care and Use Committee (AUP 16-12-033). NSG mice carry two mutations on the NOD/ShiLtJ genetic background (severe combined immune deficiency (*scid*) and a complete null allele of the IL2 receptor common gamma chain – *IL2rg*^*null*^)[36]. The *scid* mutation renders the mice B and T cell deficient whereas the *IL2rg*^*null*^ mutation leads to a deficiency in functional NK cells. The immunodeficient NOD mice share a genetic background with NSG mice and have innate immune cells deficiencies[36,37].

### Purification of recombinant α-synuclein and assembly of pre-formed fibrils

Recombinant human α-syn was purified similarly to this previously published protocol[38,39]. Briefly, the protein was expressed in BL21 *E*.*coli* transformed with a plasmid expressing human α-syn. Once expressed, cells were collected and stored at -80°C. For purification, cells were lysed by sonication and boiling, and centrifuged to remove cell debris. The α-syn-containing supernatant was dialyzed overnight in 10 mM Tris, pH 7.5, 50 mM NaCl, and 1 mM EDTA, using SnakeSkin Dialysis Tubing MWCO 7,000 (Thermo Scientific). Chromatographic separation was performed using a Superdex 200 Column (GE Healthcare Life Sciences) and a Hi-trap Q HP anion exchange column (GE Healthcare Life Sciences). Fractions containing α-syn were identified by SDS-PAGE and Coomassie staining, and then dialyzed overnight into PBS buffer (Life Sciences). A NanoDrop 2000 (Thermo Fisher) was used to determine the protein concentration by OD_280_ reading and protein was concentrated to 5 mg/mL using a Vivaspin protein concentrator spin column with a MWCO of 5kDa (GE Healthcare). Aliquots of 500 μL were stored at -80°C until use. For amyloid fibril assembly, purified recombinant α-syn was thawed and subjected to continuous shaking at 1,000 r.p.m at 37°C in a Thermomixer (Eppendorf) for 7 days. The monitoring of fibril assemblies was performed by Thioflavin T (Sigma) fluorescence reading (data not shown). Fibrils were aliquoted and frozen at -80°C until use.

### Stereotactic injections

Prior to injection, human α-syn PFFs were thawed and sonicated at RT in a water-bath cup-horn sonicator (Misonix XL2020, 50% power, 120 pulses 1 s ON, 1 s OFF for two minutes). Following sonication, we then prepared transmission electron microscope grids and stained the PFFs negatively with 1% uranyl acetate to control the morphology of the fibrils prior to injection. Grids were imaged using a Tecnai G2 Spirit TWIN transmission electron microscope at 120kV (FEI Company, Figure 1a). Mice were anesthetized with an isoflurane/oxygen mixture and injected unilaterally with either 2 μL of PFFs (5 μg/μL) or 2 μL of saline as a control in the dorsal striatum[40] (coordinates from bregma: AP: + 0.2 mm; ML: -2.0 mm; DV: - 2.6 mm from dura) at a rate of 0.2 μL/min using a glass capillary attached to a 10 μL Hamilton syringe. After injection, the capillary was left in place for 3 minutes before being slowly removed.

**Figure 1.**
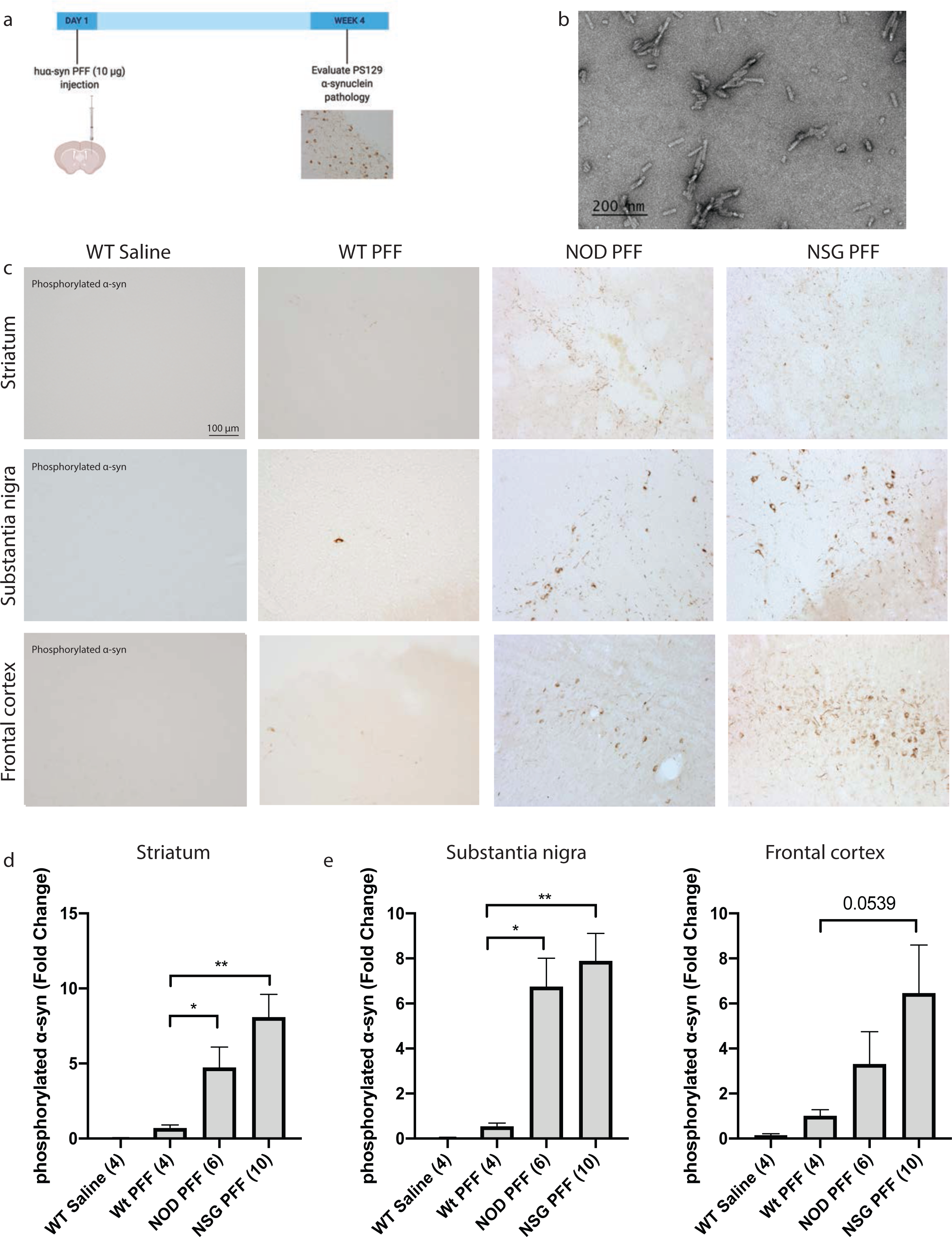
Increased phosphorylated α-syn pathology in immunocompromised mice injected with human α-syn pre-formed fibrils. a. Timeline of the experiment. b. PFFs were sonicated and validated by transmission electron microscopy. c. α-Syn pathology in the injected striatum was detected in the ipsilateral hemisphere to PFF injection in the striatum, substantia nigra and ipsilateral frontal cortex at 4 weeks post injection, by an antibody against phosphorylated α-syn at serine 129. d. Densitometry performed on 4-10 animals per group in the striatum, substantia nigra and frontal cortex (wildtype Saline, n = 4; wildtype PFFs, n = 4; NOD PFFs, n = 6; NSG PFFs n = 10). Statistical analyses were performed by Pairwise Wilcoxon Rank Sum Tests analysis* p < 0.05, ** p < 0.01, Scale bar: 100 μm.

### Adoptive transfer

Splenocytes obtained from wildtype mice were transferred i.p. (1×10^7^ cells/mouse) into NSG mice 4 weeks post PFFs injection. The optimal number of T and B cells for adoptive transfer ranges from 1×10^6^ -1×10^7^ cells/mouse[41–46]. Purification of T and B cells from total splenocytes was carried out by negative selection using Dynabeads untouched mouse T cells and Dynabeads untouched CD43 B cells isolation kits according to the manufacturer’s instruction (Invitrogen). Successful transfer of splenocytes was confirmed by flow cytometric analysis of blood and spleen at the conclusion of the study.

### Euthanasia

Mice were deeply anesthetized with sodium pentobarbital at 12 weeks post PFFs-injection. First, the spleen was rapidly collected and kept on ice and blood was collected in 0.5 mM EDTA/100 μL of blood. and kept at RT until flow cytometric analysis. Mice were then transcardially perfused with 0.9% saline followed by 4% paraformaldehyde in phosphate buffer. Brains were collected and post-fixed for 24 hours in 4% PFA, and then stored at 4°C in 30% sucrose in phosphate buffer until sectioning.

### Flow cytometric analysis

Immunophenotyping of isolated blood and splenocytes was performed. Blood was collected by cardiac puncture using a 26G needle. A minimum of 300 μL was collected into microfuge tubes containing 0.5 mM EDTA/100 μL of blood. Blood was lysed using 1x RBC lysis buffer (eBioscience), spun 300 x g for 10 mins, and the pellet was resuspended in flow buffer: PBS (minus Ca^2+^/ Mg^2+^, 2% fetal bovine serum) to bring cell number to ∼5×10^6^ cells/mL. Staining of blood cells were performed on 50 μL of blood incubated with antibodies for 30 mins, RT; SuperBright 702 anti-mouse CD45.1 (clone A20, 0.5 μg, Invitrogen) or PE-Dazzle595 anti-mouse CD45.1 (clone A20, 0.5 μg, BioLegend), monoclonal anti-mouse CD45.2 APC (clone 104, 0.125 μg, BioLegend), FITC anti mouse CD3ε (1 μg, Clone 145-2C11, BioLegend) and PE anti-mouse CD19 (clone 6D5, 0.5 μg, BioLegend). For splenocyte isolation and staining, spleens were isolated from mouse and kept on ice. In cold 1x RBC lysis buffer (eBioscience), spleens were macerated with the base of a 3 mL syringe plunger on a 70 μm cell strainer in a 10 cm petri dish on ice. 30 mL of PBS was added to stop the reaction. Cells were spun at 350 x g for 5 mins at 4°C. Pellet was resuspended in 1 mL of PBS for ∼5×10^6^ cells/mL. Staining of splenocyte cells was performed on 100 μL cells as described for blood. Following antibody incubation, blood and spleen cells were washed in flow buffer and spun at 300 x g for 10 mins. Pellets were resuspended in 300 μL of flow buffer with 1 μg/mL DAPI. Samples were acquired on a CytoFLEX S (BeckmanCoulter). Data analysis was performed using FlowJo v10.5.3. After gating on single live cells, T cells and B cells were identified in plots of CD3 vs SSC and CD19 vs SSC respectively. The mouse origin of the B and T cell susbsets was confirmed by looking at the presence of CD45.1 vs CD45.2 in each population.

### Histology

Brains were sectioned into 40 μm-thick free-floating coronal sections using a freezing microtome. Brain sections were stored in cryoprotectant and quenched with sodium peroxide. During the staining protocols, sections were incubated at room temperature overnight with primary antibodies directed against phosphorylated α-syn (rabbit anti-pS129, Abcam) at 1:10000 dilution, microglia (rabbit anti-Iba-1, WAKO) at 1:500 dilution, and an antibody to tyrosine hydroxylase (TH, rabbit, 1:1600, EMD Millipore). To detect proteinase K resistant phosphorylated α-syn, free-floating sections were incubated with proteinase K for 10 min, 10ug/mL in PBS prior to primary antibody incubation. Sections containing the substantia nigra were stained for TH and mounted onto gelatin coated slides for stereological assessment of cell counts, and counter-stained with Cresyl violet for assessment of Nissl^+^ cells. To detect CD3^+^ and MHC II in the striatum, substantia nigra and frontal cortex of mice, we used a heat induced antigen retrieval protocol using a Universal HIER antigen retrieval reagent (Abcam). Sections were incubated with rat anti-CD3 (Abcam) at 1:100 dilution or rabbit anti MHC II at 1:500 dilution (Thermo Fisher). Sections were incubated with rabbit or rat biotinylated secondary antibodies (Vector Laboratories) and conjugated with an ABC-HRP biotin/avidin complex kit (Vector Laboratories). Staining was developed with 3,3′-diaminobenzidine then sections were mounted for imaging and analysis. Representative images of phosphorylated α-syn pathology and microglia were acquired at 20x magnification using a Nikon Eclipse Ni-U microscope.

### Immunofluorescence

To detect CD4^+^ cells we used heat induced antigen retrieval (as detailed above). Sections were incubated with rabbit anti-CD4 (Abcam) at 1:100 dilution. Next, sections were treated with donkey anti-rabbit Alexa 594 (1:500 dilution, Invitrogen) with DAPI (1:10000 dilution) added to the secondary solution. Sections were mounted and coverslipped using Vectashield mounting medium (Vector Laboratories) Representative images of CD4^+^ were acquired at 20x magnification using a Nikon Eclipse Ni-U microscope.

### Image J analysis of phosphorylated α-syn pathology, MHC II and CD4^+^ immunofluorescence

We investigated phosphorylated α-syn pathology in the ipsilateral hemisphere to the PFF injection and in the contralateral striatum, substantia nigra and frontal cortex of mice injected with human α-syn PFFs by densitometry. Briefly, we acquired photomicrographs from slides at 20x magnification on three consecutive sections, three images per section, distanced by a 240 µm interval. Striatal images, were captured at the level of bregma +0.26 mm, the images from the substantia nigra were captured from bregma -3.08 mm and the frontal cortex images from bregma +3.20 mm. The images of phosphorylated α-syn and MHC II stained tissue were then analyzed in ImageJ64 (Rasband WS (1997) ImageJ (modified in 2016) NIH, Bethesda, Maryland, USA). Images were converted to 8-bit grayscale. We set thresholds for each image and analyzed particles in order to obtain the size of the area and the mean grey value (A.U.) of the phosphorylated α-syn-positive regions. We determined the average grey value in each brain region and animal. We then normalized to the PFFs injected wildtype mice (value as 1) and plotted the groups as a fold change from the PFFs injected wildtype group. For CD4^+^ immunofluorescence, 20x images were analyzed by Image J. Images were converted to 8-bit grayscale. We set a threshold for each section using the triangle setting, and then converted images to black background (of binary masks). We recorded the fraction of each area (striatum, substantia nigra and frontal cortex) that was positive and expressed it as a percentage.

### Assessment of microglial morphology

Microglia morphology was assessed as previously described[47]. Color (RGB) images of Iba-1-stained striatum, substantia nigra and frontal cortex tissues were acquired bilaterally at 60x magnification (oil immersion 1.40 N.A.) using a Nikon Eclipse Ni-U microscope. A total of nine images/animal were analyzed; three images from three sequential brain sections through the striatum (as described above). To assess the morphology of microglia in the samples, RGB color images were processed by a custom MATLAB (Mathworks) script. Cell bodies and other large regions were segmented first. Dynamic thresholds were determined for both the blue intensity and saturation channels of the image. Upper and lower bounds of the thresholds were set to match the full width at half maximum of curves fit to the histograms of each channel. If the signal met the required criteria for either the blue channel or the saturation channel thresholds, it was selected for further analysis. Segmented regions were filtered to remove small (under 2,000-pixel) areas and to remove objects that touched the borders of the image. The segmented areas were eroded to more accurately conform to cell bodies. The ratio of area:perimeter (referred to as hydraulic radius) was calculated and used as a measure for microglial activation; activated microglia are amoeboid in shape and therefore exhibit a larger index score.

### Stereological counting

We used a computer-assisted cell quantification program (StereoInvestigator, MBF Bioscience) coupled to a Nikon Eclipse Ni-U microscope (Nikon). In each mouse (3–4 animals per group), we analyzed 5–7 sections containing substantia nigra. They were spaced by 240 μm (section interval = 6). Contours of the region were drawn at 10x magnification (air immersion, N.A. 0.45). Quantifications were performed at 60x (oil immersion, N.A. 1.40) using a counting frame of 100 μm × 100 μm, grid size set to 200 × 200 μm, with a guard zone of 2 μm, and dissector height set at 12 μm.

### Statistical analysis

All values are expressed as mean ± SEM. Differences in means between the groups were analyzed using a Pairwise Wilcoxon Rank Sum Tests or one-way ANOVA test by using R software (v 3.6.2) and GraphPad Prism software, respectively. A *p* value < 0.05 was considered statistically significant.

## RESULTS

### Increased phosphorylated α-syn inclusions in NSG PFFs injected mice

To determine the influence of a compromised immune system in the accumulation of phosphorylated α-syn following injection of fibrillar α-syn into the striatum, we compared NSG mice to NOD/ShiLtJ and to wildtype mice[37]. The timeline of the experiment is represented in Figure 1a. The sizes of the PFFs were verified by transmission electron microscopy (Figure 1b), which were consistent with the PFFs used in our previous studies[39].

The accumulation of phosphorylated α-syn inclusions in wildtype, NOD and NSG mice four weeks after a single intrastriatal injection with recombinant human α-syn PFFs are presented in Figure 1. Pathology in three brain regions (ipsilateral to the PFF injection), i.e. the striatum, substantia nigra and frontal association cortex (referred to as frontal cortex from hereon, see Figure 1c for representative images of pathology), was selected for quantification. In all graphs, the levels of phosphorylated α-syn were normalized to the average level of pathology in the wildtype mice injected with α-syn PFFs (this is set to 1) and plotted as a fold change.

In the striatum and substantia nigra, NSG mice inoculated with α-syn PFFs displayed significantly more pathology compared to wildtype mice injected with α-syn PFFs. In the frontal cortex, the levels of phosphorylated α-syn pathology showed a non-significant trend for an increase in PFF-injected NSG mice compared to wildtype PFFs-injected mice (p = 0.0539, Figure 1d).

We observed phosphorylated α-syn pathology in PFFs injected NOD mice (Figure 1c and 1d) that was not different to that seen in NSG mice (p >0.05, Figure 1c and 1d). NOD/ShiLtJ mice have multiple immune cell dysfunctions including defects in innate immunity, like reduced dendritic cell function, lack of mature NK cells and defective macrophage activity, as well as neuroinflammatory changes[36,37,48]. They also spontaneously develop type I diabetes (43%-80% by 30 weeks of age)[37,49–51]. The phosphorylated α-syn load in the NOD/ ShiLtJ mice was not significantly different to that which we observed in NSG PFFs-injected mice, which supports the hypothesis that immune defects can lead to increased phosphorylated α-syn pathology. To avoid the potential complication of NOD/ShiLtJ mice developing type I diabetes, we injected NSG mice with PFFs for the remainder of the study, which aimed to address whether T or B cells influence the accumulation of phosphorylated α-syn inclusions. We used wildtype mice as the donors of the T and B cells for the remainder of the study.

### Decreased phosphorylated α-syn in NSG mice reconstituted with T cells

In the results described above, we established that there is increased α-syn related pathology following intrastriatal PFFs injection in NSG mice compared to wildtype mice. In our next set of experiments, we investigated whether reconstituting the T cell population in PFFs-injected NSG mice would alter the levels of phosphorylated α-syn in the striatum, substantia nigra or frontal cortex. Throughout the manuscript, we refer to NSG mice injected with human α-syn PFFs and reconstituted with T cells as NSG PFF T and NSG mice injected with human α-syn PFFs and reconstituted with B cells as NSG PFF B. We conducted the reconstitution of T cells in two separate experiments and the results were pooled after we determined that there was no difference in the neuropathological outcomes between the two experiments (data not shown). Mice were injected with PFFs in the striatum; four weeks later, the adoptive transfer of immune cells was performed. Pathology was assessed 12 weeks post PFFs injection. Flow cytometry plots from the blood and splenocytes isolated from the reconstituted NSG mice were positive for CD3^+^ CD45.2^+^ T cells. C57Bl/6J mice express the CD45.2 allele while NSG mice express the CD45.1 allele (representative plots in Figure 2a), therefore CD45.2 positive cells demonstrated the successful uptake of donated T cells. The gating strategy is provided in supplementary Figure 1. Supplementary Tables 1 and 2 contain individual counts of T cells/µL in blood and spleen flow samples (Tables 1 and 2, respectively). We observed CD3^+^ cells in the striatum, substantia nigra and frontal cortex in PFFs injected mice that were reconstituted with T cells (Figure 2b). As CD3 is also expressed on NK cells, we confirmed the presence of CD4^+^ T cells in the striatum, substantia nigra and frontal cortex by immunofluorescence and quantified the CD4^+^ signal in the striatum, substantia nigra and frontal cortex collectively (Figures 2c, d).

**Figure 2.**
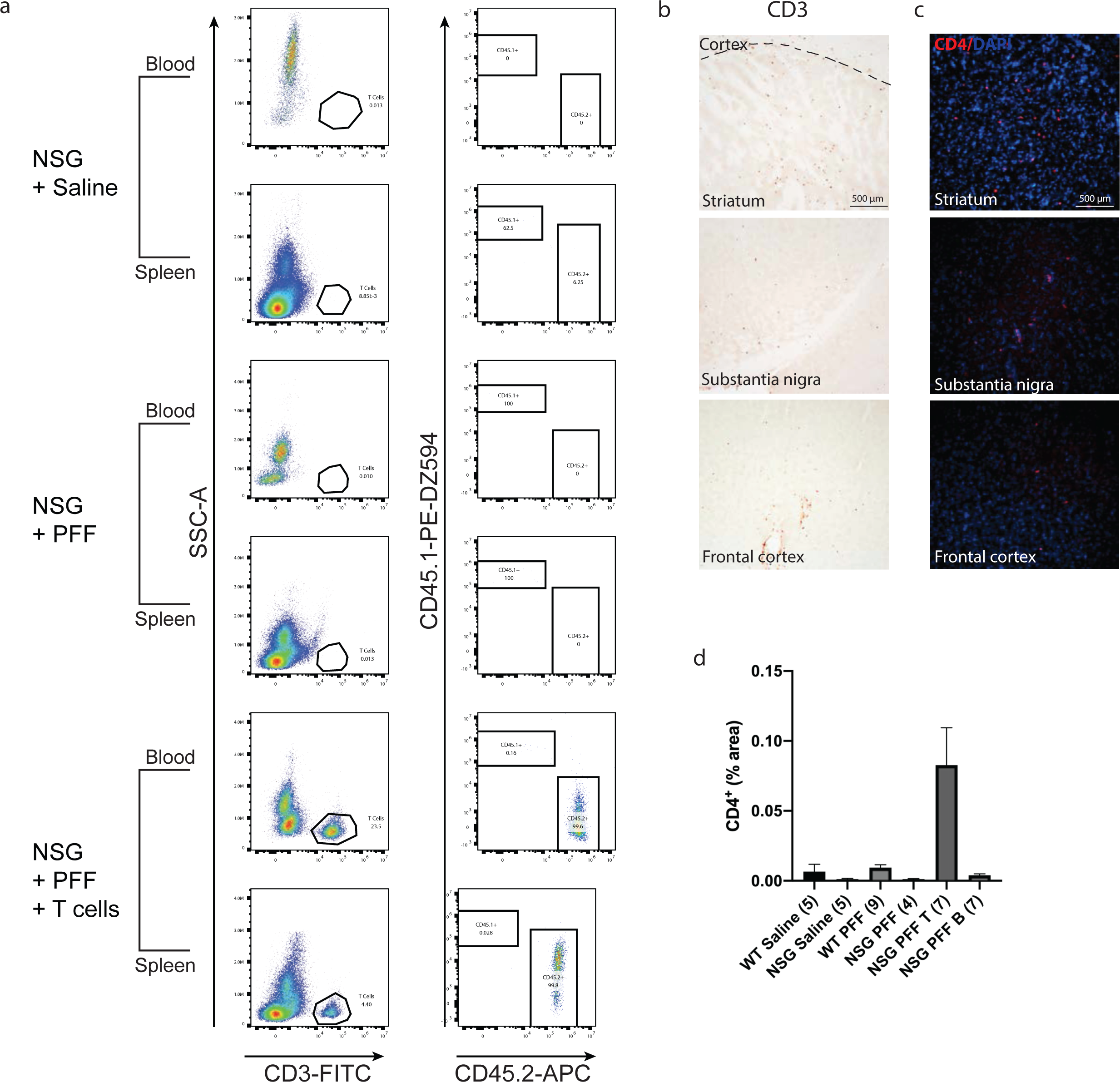
T cells in the blood, spleen and brain following adoptive transfer. a. Flow cytometric analysis. Wildtype mice contained populations of T and B cells that are CD45.2 positive. NSG mice did not contain T and B cell populations. Following adoptive transfer of T cells to NSG mice, CD45.2^+^ CD3^+^ T cells were detected. Representative plots are shown for each treatment condition. b. Representative images from the mouse striatum, substantia nigra and frontal cortex staining positive for CD3^+^T cells in mice that received adoptive transfer of T cells. c. Immunofluorescent staining for CD4^+^ T cells in the striatum, substantia nigra and frontal cortex. d. Percentage area of tissue that is positive for CD4 signal in striatum, substantia nigra and frontal cortex combined. Scale bar: 100 μm.

The PFFs-injected NSG mice exhibited clearly increased phosphorylated α-syn pathology compared to PFFs-injected wildtype mice in the substantia nigra and frontal cortex and not in the striatum (Figure 3b, c, *p* < 0.05). Because the phosphorylated α-syn pathology in the contralateral hemisphere (Supplementary Figure 2 a, b) was not as robust as the ipsilateral side within the experimental timeline (Figure 3a), which would make it difficult to evaluate the changes caused by reconstituted T or B cells, we focused our analyses on the ipsilateral hemisphere. Notably, reconstituting T cells in NSG PFFs injected mice significantly decreased levels of phosphorylated α-syn in the substantia nigra when comparing NSG PFFs to NSG PFF T mice (Figure 3b, c, *p* < 0.05). The levels of phosphorylated α-syn in the striatum and frontal cortex following T cell reconstitution decreased, but did not reach significance (Figure 3c, *p* > 0.05).

**Figure 3.**
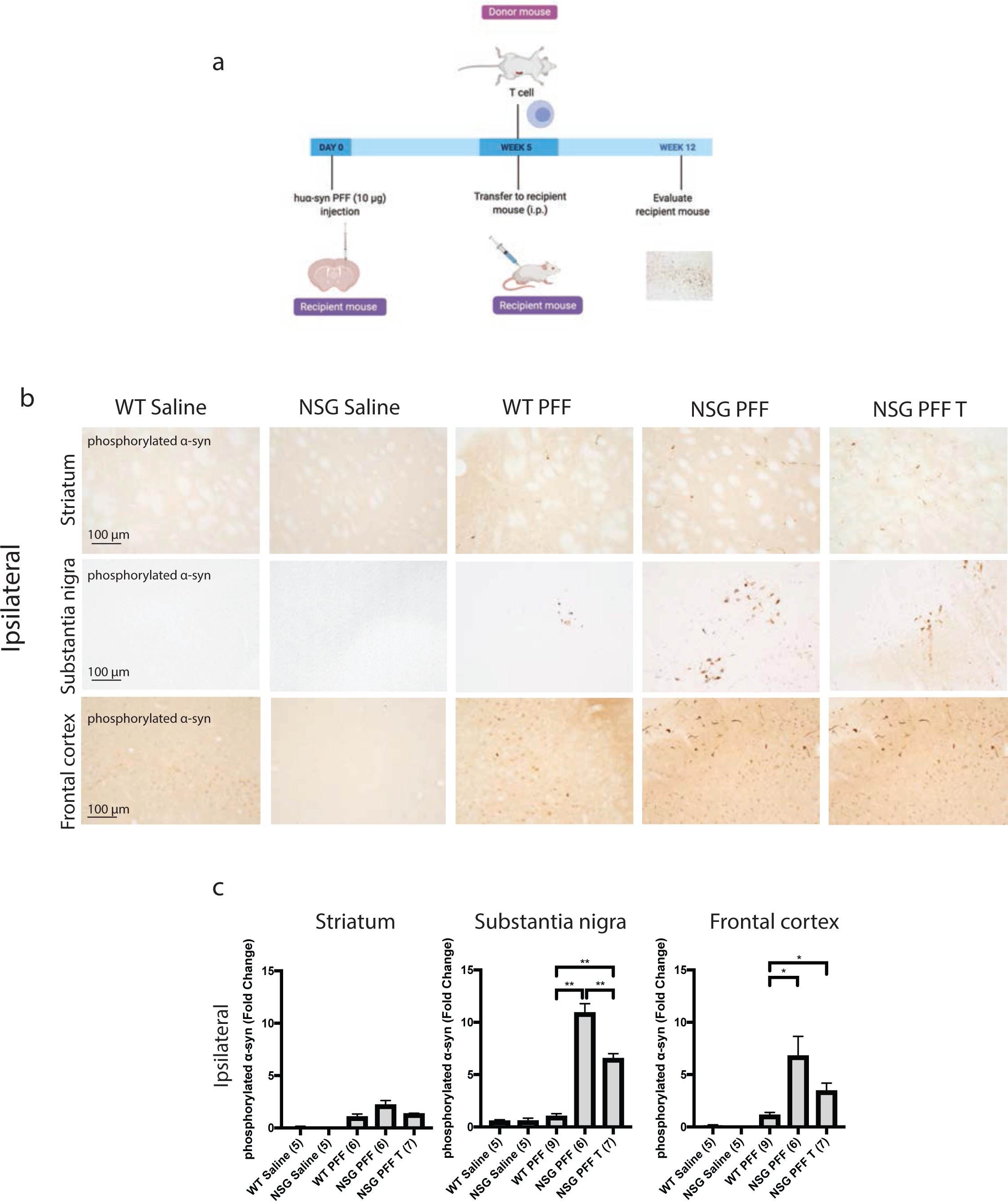
Reduced phosphorylated α-syn pathology in immunocompromised mice that received adoptive transfer of T cells. a. Timeline of experiment. b. Phosphorylated α-syn was detected in the ipsilateral hemisphere to PFF injection in the striatum, substantia nigra and frontal cortex. The reconstitution of T cells was conducted in two separate experiments and the results pooled after results of the reduction in phosphorylated α-syn were shown to be consistent between the two experiments. c. Densitometry of 5-9 mice per group to determine the fold change in phosphorylated α-syn levels in the ipsilateral striatum, substantia nigra and frontal cortex. Wildtype Saline, n=5; NSG Saline n=5, wildtype PFF, n=6; NSG PFF, n=6; NSG PFF T n=7). d. Statistical analyses were performed by Pairwise Wilcoxon Rank Sum Tests analysis * p<0.05, ** p<0.01. Scale bar: 100 μm.

As a control experiment for the reconstitution of T cells, we reconstituted a different group of NSG mice with B cells (timeline of experiment represented in Figure 4a). Since B cells on their own are not expected to affect pathology as they need T cells to work in concert for an antibody response[52,53], we used this experiment as a control to determine the effect of the reconstitution per se. The transfer of B cells was confirmed by flow cytometry (Figure 4b, supplementary Tables 3 and 4). Consistent with the above results, there was an increase in phosphorylated α-syn in the ipsilateral substantia nigra in PFFs-injected NSG mice compared to PFFs-injected wildtype mice (*p* < 0.05, Figure 4c, d). There was a significant increase in phosphorylated α-syn pathology in the ipsilateral striatum and substantia nigra in the wildtype PFFs-injected mice compared to NSG PFF B (Figure 4c, d, *p* < 0.05). In the ipsilateral striatum, substantia nigra and frontal cortex, the levels of phosphorylated α-syn pathology in the PFFs-injected NSG mice following the transfer of B cells did not significantly change (Figure 4c, d). The levels of phosphorylated α-syn in the contralateral hemisphere did not significantly differ between the groups (supplementary Figure 2 c, d).

**Figure 4.**
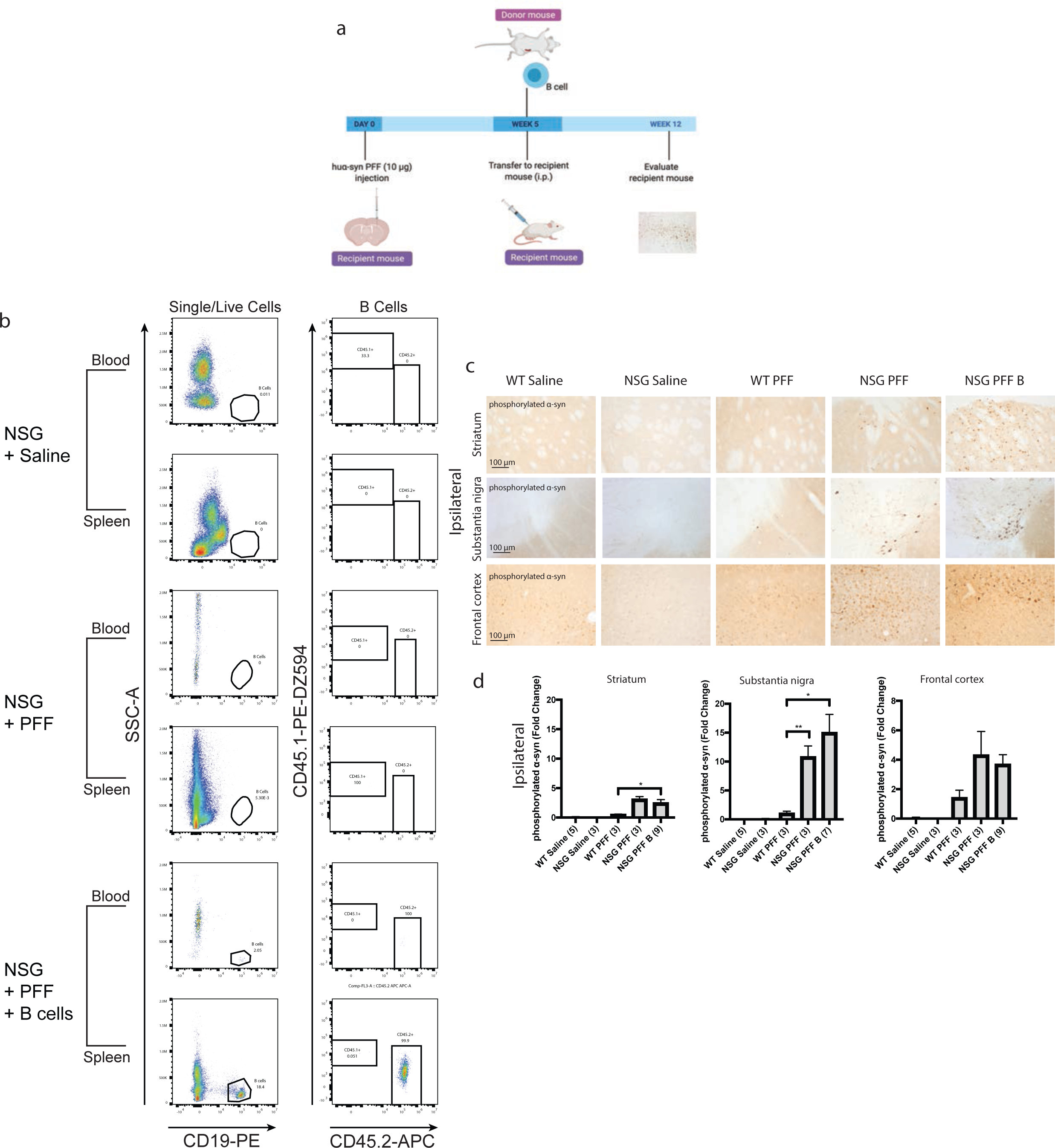
Adoptive transfer of B cells alone did not alter phosphorylated α-syn pathology in immunocompromised mice. a. Timeline of experiment. b. Flow cytometric analysis of mouse spleen and blood following adoptive transfer demonstrated that wildtype mice contain populations of T and B cells that are CD3 and CD19 positive. NSG mice do not have T and B cell populations. Following adoptive transfer of B cells, NSG mice contained CD45.2^+^ CD19^+^ B cells. Representative plots are shown for each treatment condition. c. Phosphorylated α-syn was detected in the ipsilateral striatum, substantia nigra and frontal cortex. d. Densitometry of 3-9 mice per group to determine the fold change in phosphorylated α-syn levels in the ipsilateral striatum, substantia nigra and frontal cortex. Wildtype Saline, n = 5; NSG Saline n = 3, wildtype PFFs, n = 3; NSG PFFs, n = 3; NSG PFF B n = 9). The error bars represent S.E.M. Statistical analyses were performed by Pairwise Wilcoxon Rank Sum Tests analysis ** p < 0.01. Scale bar: 100 μm.

We also evaluated the presence of proteinase K resistant phosphorylated α-syn as a readout for aggregated α-syn[54,55]. In all groups injected with PFFs, we observed proteinase K-resistant phosphorylated α-syn (supplementary Figure 3). Proteinase K-resistant phosphorylated α-syn was also present in the striatum, substantia nigra and frontal cortex.

### Microgliosis in mice reconstituted with T cells

To understand whether microglia were involved in the reduction of phosphorylated α-syn pathology in NSG mice reconstituted with T cells, we analyzed the morphology of microglia. We immunostained for Iba-1 to compare microglial morphology between all groups (Figure 5). We compared the morphology of Iba-1 immunoreactive microglia from the same anatomical level of the ipsilateral and contralateral striatum, substantia nigra and frontal cortex in mice from each experimental group. Specifically, we used a MATLAB script to define the ratio between the perimeter and surface area of Iba-1-immunoreactive microglia, allowing us to calculate the hydraulic radius of each cell as an index of the activation state as previously[47]. The area:perimeter index (hydraulic radius) measures microglial activation as activated microglia are amoeboid in shape. Thus, these activated cells have a large area and small perimeter, increasing the index score. The hydraulic radius was significantly increased in microglia from within the striatum of NSG PFFs injected mice that received T cells relative to wildtype Saline, wildtype PFFs, NSG Saline, NSG PFFs, NSG PFF T and NSG PFF B cell injected mice (Figure 5b, *p* < 0.05) indicating that microglia in the NSG PFF T group are activated. The hydraulic radius of microglia analyzed from the substantia nigra was not significantly different between the groups (Figure 5b, *p* > 0.05). In the frontal cortex, the hydraulic radius of microglia in the NSG PFF B mice was significantly reduced compared to the NSG PFF T mice (Figure 5b, *p* < 0.05). Microglia morphology in the contralateral striatum, substantia nigra and frontal cortex did not significantly change between groups (Figure 5b, p>0.05).

**Figure 5.**
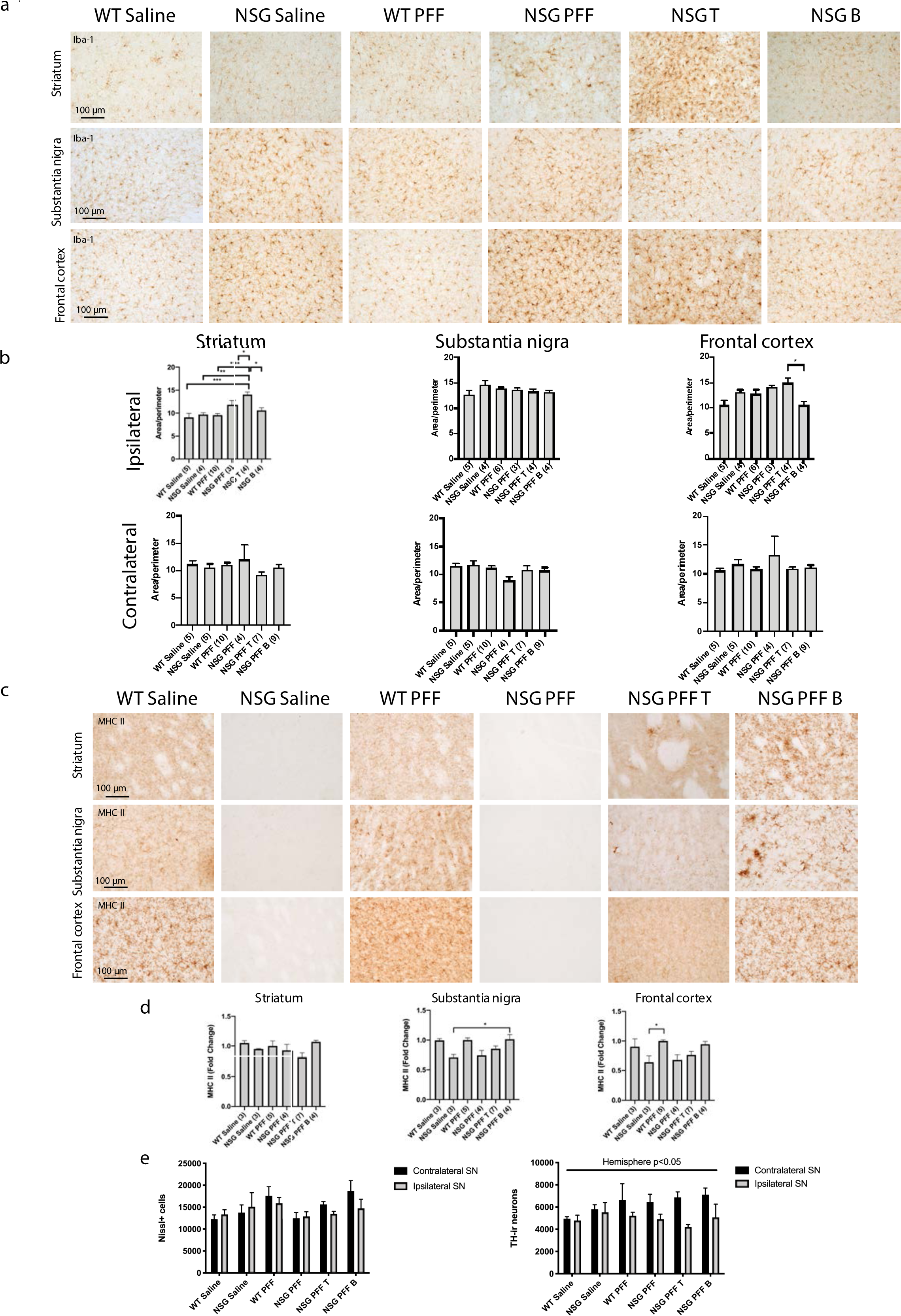
Microgliosis and dopaminergic cells death in the brain of α-syn PFFs injected mice. a. Representative images of Iba-1 immunoreactive microglia were present in the ipsilateral striatum, substantia nigra and frontal cortex of saline and PFFs injected mice. b. Quantification of microglia morphology in the ipsilateral hemisphere to PFF injection and in the contralateral striatum, substantia nigra and frontal cortex (area/perimeter). Wildtype Saline, n = 5; NSG Saline n = 4, wildtype PFFs, n = 10; NSG PFFs, n = 3; NSG PFF T n = 4; NSG PFF B n = 4). c. Representative images of MHC II immunoreactive cells in the ipsilateral striatum, substantia nigra and frontal cortex of saline and PFFs injected mice. d. Densitometry of 3-7 mice per group to determine MHCII levels in the ipsilateral substantia nigra. Wildtype Saline, n = 3; NSG Saline n = 3, wildtype PFFs, n = 5; NSG PFFs, n = 4; NSG PFF T n = 7; PFF B n = 4). e. Stereological counts of saline and PFFs injected wildtype and NSG mice. There was no main effect of genotype (*p* > 0.05), but a significant main effect of hemisphere (*p* < 0.05) in the total number of TH-expressing cells in the substantia nigra between contralateral substantia nigra (black bars) and ipsilateral substantia nigra (grey bars). The error bars represent S.E.M. Statistical analyses were performed by one and two-ANOVA analysis * p <0.05, ** p < 0.01, *** p < 0.001. Scale bar: 100 μm.

Several single nucleotide polymorphisms associated with PD risk are found in and around the human leukocyte antigen (HLA) locus coding for MHC II[56]. MHC II is used by antigen presenting cells (e.g. microglia) to interact with T cells. We evaluated the levels of MHC II by immunohistochemistry on sections through the striatum, substantia nigra and frontal cortex (Figure 5d). There was an absence of signal in the NSG Saline and NSG PFF mice. There were no significant changes in the MHC II signal in the striatum (Figure 5d, p> 0.05). In the substantia nigra, MHC II signal was significantly higher in the NSG PFF B group compared to NSG Saline (Figure 5d, p< 0.05). In the frontal cortex, there was a significant increase in the MHC II signal in the wildtype PFF group compared to the NSG Saline group (Figure 5d, p< 0.05).

To determine whether adoptive transfer of T cells altered the number of surviving dopamine neurons in the substantia nigra, we performed stereological cell counts on TH-immunostained sections from wildtype saline, wildtype PFFs-injected and NSG saline and NSG PFFs-injected mice and from NSG mice reconstituted with T and B cells. We determined that PFFs injection resulted in significant cell loss in TH-positive neurons (Figure 5d, significant main effect of hemisphere, *p* <0.05) and that reconstitution of T or B cells was not significantly different from the cell loss observed in the NSG PFFs-injected mice (Figure 5d, *p* >0.05).

## DISCUSSION

We explored the role of T cells in the accumulation of α-syn following intracerebral inoculation with α-syn fibrils in immunocompromised mice (NSG) that lack B, T and natural killer cells[36]. Compared to wildtype mice injected with α-syn fibrils into the striatum, NSG mice injected with PFFs injections developed greater accumulation of phosphorylated α-syn in the striatum, substantia nigra and frontal cortex. To identify whether the absence of T lymphocytes was driving this elevation in phosphorylated α-syn inclusions, we reconstituted NSG mice with T cells via adoptive transfer four weeks after intrastriatal injections of α-syn PFFs. To control for the injection of immune cells via adoptive transfer, we injected a separate group of mice with B cells isolated from wildtype mice. Notably, the transfer of T cells resulted in a significant decrease in phosphorylated α-syn immunostaining in the substantia nigra, but not in the striatum and frontal cortex, where only a trend was observed. Our data imply that a critical number of T cells are required to alter the presence of α-syn pathology *in vivo*, as with the adoptive transfer, significantly more T cells were found in the brain than in PFFs injected mice. Such an increase in the number of infiltrating T cells could mediate the decrease in phosphorylated α-syn pathology in the substantia nigra. This complements several reports associating altered T cell function with PD and MSA[21–26,57] and studies demonstrating activation of immune cells by pathogenic α-syn[27,31,58]. One such study measuring nigral neurodegeneration as the readout supports the hypothesis that immune cells can modulate neuroinflammation and neurodegeneration in PD mouse models[59]. Overexpression of human α-syn in the substantia nigra of immune competent mice results in microglial activation, MHC II activation, T cell and monocyte entry into the brain and consequently, loss of substantia nigra neurons[33,60,61]. Our findings support these prior studies by showing that intrastriatal injection of human α-syn PFFs leads to elevated phosphorylated α-syn pathology in immunodeficient compared to wildtype mice[62]. The exacerbation of pathology observed in PFFs injected immunocompromised mice was partially mitigated by adoptive transfer of T cells. Further, following adoptive transfer we observed CD4^+^ T cells invading the striatum, substantia nigra and frontal cortex, which supports previous work showing infiltration of T cells in PD brains and in a neurotoxin model of nigral neurodegeneration[19,20]. We observed CD3^+^ CD4^+^ cells throughout the structures we studied, however due to the small number of infiltrating cells, it was difficult to determine absolute numbers and their distribution. The regional distribution of the cells in the brain may account for why T cells were more effective in reducing nigral pathology, but not striatal or cortical pathology. Although we could not determine regional differences in T cell infiltration in the brain, our findings of CD3^+^ and CD4^+^ T cells in the brain align well with a growing body of evidence that T cells invade the central nervous system in neurodegenerative conditions[29,63,64].

We also investigated whether the decrease in pathology observed following adoptive transfer of T cells could be due to cytotoxic T cells killing neurons that exhibit phosphorylated α-syn, leading to an apparent reduction of α-syn staining as a consequence. This idea would be consistent with an earlier study showing that mice lacking CD4^+^ T cells are relatively resistant to MPTP-induced degeneration of substantia nigra dopamine neurons[19]. However, we did not find this to be the explanation for the reduction in α-syn pathology. Stereological counts of dopaminergic neurons of the substantia nigra revealed similar reductions in nigral TH-immunoreactive neurons following intrastriatal injection of α-syn PFFs in mice lacking T cells as those having their T cell population reconstituted (Figure 5d).

An alternative mechanism for the significant decrease in phosphorylated α-syn following T cell reconstitution is that the infiltration of T cells leads to the activation of microglia and that these resident macrophages are involved in removing phosphorylated α-syn. Earlier studies have also shown that T cells invading the brain can activate microglia[63–65]. Indeed, we also found microglia morphology was significantly altered in the striatum of mice that received adoptive transfer of T cells following intrastriatal injection of α-syn PFFs. The microgliosis that we observed was restricted to the striatum, in line with previous work in immunocompetent rats[66]. Microglia are heterogenous and have distinct region-dependent transcriptional identities, probably because the local environment influences their molecular and morphological profiles[67–69]. It is therefore possible that the response to human α-syn is region-specific, with striatal microglia becoming more highly activated than microglia in the substantia nigra and frontal cortex in this paradigm. We also observed no increase in MHCII signal in PFF-injected NSG mice that received an adoptive transfer of T cells. This is in contrast to what has been reported recently, in where T cells are necessary for the upregulation of MHC II in microglia, which then leads to loss of dopamine neurons in the substantia nigra in response to viral overexpression of α-syn in rats[34]. The absence of MHC II upregulation in response to α-syn PFFs (in contrast to the model with viral overexpression) could be due to recombinant adeno-associated virus being inflammatory itself[67–70]. Alternatively, the α-syn species generated by the recombinant adeno-associated virus compared to PFFs are different. In contrast to PFFs, vector overexpressing models do not efficiently generate stable, seeding assemblies, and the toxic effects are believed to be mediated via overexpression of soluble α-syn[71]. Therefore the α-syn that is generated by models using recombinant adeno-associated virus and PFFs can stimulate the immune system differently.

Our study did not address the influence of NK cells in the accumulation of phosphorylated α-syn inclusions. NK cell function is altered in peripheral lymphocytes obtained from PD patients as the levels of inhibitory receptors on NK cells are significantly lowered in PD patients, rendering the NK cells more susceptible to activation[72]. NK cells are one of the first lines of defense of the innate immune system[73]. They respond rapidly to a variety of insults with cytolytic activity and cytokine secretion[74]. They are implicated in diseases of autoimmunity and within the CNS, and can interact with microglia[74]. Recently, it was reported that systemic depletion of NK cells in a preclinical model of PD exacerbated α-syn pathology[75]. That particular preclinical model utilized α-syn transgenic mice that over-express human α-syn with the A53T mutation, and combined it with an injection of human α-syn PFFs into the dorsal striatum. The depletion of NK cells in this model of α-syn-induced pathology suggests that NK cells can contribute to the clearance of α-syn aggregates, or prevent their formation[75]. The mice we used in our study also have deficiencies in NK cells. Mice that have a mutation in the interleukin-2 receptor common gamma chain, such as the NSG mice used in this study, have interrupted cytokine signaling networks for multiple cytokines, in particular Interleukin-15, which contributes to a complete lack of NK cells[76]. Thus, the NOD/ShiLtJ mice have dysfunctional NK cells and NSG mice lack NK cells all together[36,37,76]. Notably, we found that phosphorylated α-syn was increased in multiple brain regions in both NOD/ShitLtJ mice and NSG mice when compared to wildtype mice in Figure 1. In light of the findings mentioned above[75], a driver of this difference between wildtype and the mice on the NOD background could also be the dysfunction or absence of NK cells. Thus, we can be speculate that both arms of immunity are involved in modulating accumulation of phosphorylated α-syn. To clarify the specific role of NK cells in α-synucleinopathy, future studies are needed where NK cells are reconstituted in the immunocompromised mice.

While our results highlight the role for the adaptive immune system, and specifically T cells, in the accumulation of α-syn, we did not investigate the dynamics of the microglial response and accumulation of phosphorylated α-syn. To further delineate the role for T and B cells in the accumulation of α-syn in a paradigm relevant to PD, experiments transferring T and B cells from wild type PFFs-injected mice that have been previously exposed to human α-syn PPFs would be informative. This would also allow us to define the role of B cells, as B cells require T cells for their maturation and for the production of antibodies[53]. In the current study, we injected naïve B-cells from wild-type mice that had never been exposed to human α-syn PFFs, therefore the B cells lack the specific receptors for different antigens present in the mixture of α-syn assemblies that PFFs contain to mount a mature antibody response.

## CONCLUSION

Compared to wildtype mice injected with PFFs into the striatum, NSG immunocompromised mice injected with PFFs exhibited increased phosphorylated α-syn levels in the substantia nigra, and not in the striatum and frontal cortex. The accumulation of phosphorylated α-syn in the substantia nigra was reduced when we used adoptive transfer to reconstitute the T cell population in the immunocompromised mice. We observed signs of elevated microglia activation in the striatum in mice that received the T cell adoptive transfer. Taken together, our study provides direct *in vivo* evidence that T cells modulate accumulation of phosphorylated α-syn and supports an important role for the immune system in PD pathogenesis.

## Supporting information

All supplemental files

## List of abbreviations

MHC II: major histocompatibility complex II
NOD/ShiLtJ (NOD): Non-obese diabetic
NSG: NOD Scid gamma
PD: Parkinson’s disease
PFFs: preformed fibrils
α-syn: α-synuclein

## Acknowledgements

We thank Dr. Corinne Esquibel from the Optical Imaging core for developing the MATLAB algorithm for microglia morphology analysis, Emily Wolfrum from the Bioinformatics and Biostatistics for assistance with the statistical analysis and the staff of the Vivarium of Van Andel Research Institute for animal care. We acknowledge the Van Andel Institute and the many individuals and corporations that supported financially the neurodegenerative research at Van Andel Research Institute. S.G. was supported by the Peter C. and Emajean Cook Foundation and the Farmer Family Foundation. Research reported in this publication was supported by the National Institute On Deafness And Other Communication Disorders of the National Institutes of Health under Award Numbers R01DC016519 (P.B.) and the National Institute of Neurological Disorders and Stroke of the National Institutes of Health under Award Numbers R21NS106078-01A1 (P.B.). The content is solely the responsibility of the authors and does not necessarily represent the official views of the National Institutes of Health. We would like to acknowledge iPark and the Farmer Family Foundation for funding a portion of this work.

## Funding

This work was supported by the Farmer Family Foundation and S.G. was supported by the Peter C. and Emajean Cook Foundation.

## Conflict of interest

PB has received commercial support as a consultant from Axial Biotherapeutics, Calico, CuraSen, Fujifilm-Cellular Dynamics International, Idorsia, IOS Press Partners, LifeSci Capital LLC, Lundbeck A/S and Living Cell Technologies LTD. He has received commercial support for grants/research from Lundbeck A/S and Roche. He has ownership interests in Acousort AB and Axial Biotherapeutics and is on the steering committee of the NILO-PD trial. The authors declare no additional competing financial interests.

## Authors’ contributions

SG designed the study, performed the surgeries, adoptive transfer, prepared samples for flow cytometry, the immunohistochemistry, acquisition of brightfield images, and MATLAB and image analysis, interpreted the histological results, and wrote the manuscript. TT contributed to the design of the study, assisted in the isolation of cells for adoptive transfer and edited the manuscript. NLR assisted in the NOD and NSG PFFs surgeries and edited the manuscript. RS acquired and analyzed the flow cytometry data. WP performed the Iba-1 and CD4 immunostaining. KB generated the PFFs. ES and LM assisted with perfusions and performed immunohistochemistry. ARB assisted with the adoptive transfer. MLEG, JAS, and LB edited the manuscript and contributed to interpretation of the results. AP contributed to interpretation of the results. JM provided the PFFs and edited the manuscript. PB contributed to the design of the study and interpretation of the results and edited the manuscript. All authors gave input to the manuscript. All authors read and approved the final manuscript. VL passed away when this manuscript was being finalized.

## Supplementary Material

**Supplementary Figure 1. Flow cytometry gating strategy**. Live single cells were gated for T cells based on CD3 or B cells using CD19. The WT-mouse origin of cells was confirmed by the presence of CD45.2. NSG-derived cells would express CD45.1.

**Supplementary Figure 2. Phosphorylated α-syn is found in the contralateral hemisphere in PFFs-injected mice**. a. Tissue from the contralateral hemisphere of the striatum, substantia nigra and frontal cortex from wildtype saline, NSG saline, wildtype PFF, NSG PFF and NSG PFF T mice were all positive for phosphorylated α-syn. b. Densitometry of 5-7 mice per group to determine the fold change in phosphorylated α-syn levels in the contralateral striatum, substantia nigra and frontal cortex. Wildtype Saline, n = 5; NSG Saline n = 5, wildtype PFFs, n = 9; NSG PFFs, n = 4; NSG PFF T n = 7). c. Tissue from the contralateral hemisphere of the striatum, substantia nigra and frontal cortex from wildtype saline, NSG saline, wildtype PFF, NSG PFF and NSG PFF B mice were all positive for phosphorylated α-syn. d. Densitometry of 5-7 mice per group to determine the fold change in phosphorylated α-syn levels in the contralateral striatum, substantia nigra and frontal cortex. Wildtype Saline, n = 5; NSG Saline n = 5, wildtype PFFs, n = 9; NSG PFFs, n = 4; NSG PFF B n = 7). The error bars represent S.E.M. Statistical analyses were performed by Pairwise Wilcoxon Rank Sum Tests analysis ** p < 0.01. Scale bar: 100 μm.

**Supplementary Figure 3. Proteinase K resistant phosphorylated α-syn is found in PFFs-injected mice**. Tissue from the striatum, substantia nigra and frontal cortex from wildtype saline, NSG saline, wildtype PFF, NSG PFF, NSG PFF T and NSG PFF B mice were all positive for phosphorylated α-syn following proteinase K treatment. Scale bar: 100 μm.

**Supplementary Table 1. 45.2**^**+**^ **B cells/μL in flow blood sample**

**Supplementary Table 2-45.2**^**+**^ **B cells/μL in flow spleen sample**

**Supplementary Table 3-45.2**^**+**^ **T cells/μL in flow blood sample**

**Supplementary Table 4-45.2**^**+**^ **T cells/μL in flow spleen sample**

## Notes

### Competing Interest Statement

The authors have declared no competing interest.

### Summary of Updates

We have added a reference to further support the statement that "Viral overexpression of human alpha-syn in the substantia nigra of immune competent mouse results in microglial activation, MHC II activation, T cell and monocyte entry into the brain and consequently, loss of substantia nigra neurons" on page 12. we have made numerous edits to the discussion and We have added data to quantify the phosphorylated alpha-syn and microgliosis in the contralateral hemisphere, quantify the CD4+ cells in the brain, quantify MHCII in the ipsilateral hemisphere, new gating strategy for flow cytometry data and stained for proteinase-K resistant phosphoryalted alpha-syn.

