## Supplementary material for "T cells limit accumulation of aggregate pathology following intrastriatal injection of α-synuclein fibrils": All supplemental files

### Supplementary Tables 1-4

Table 1- CD45.2$^+$ B cells/μL in flow blood sample

| Wt Saline | NSG Saline | Wt PFF | NSG PFF | NSG PFF T | NSG PFF B |
|---|---|---|---|---|---|
| 11.4 | 0.0 | 0.0 | 0.0 | 0.0 | 0.0 |
| 138.6 | 0.0 | 0.0 | 0.0 | 0.0 | 0.0 |
| 191.7 | 0.0 | 0.0 | 0.0 | 0.1 | 0.3 |
| 32.4 | 0.0 | 0.0 | 0.0 | 0.4 | 0.2 |
| 48.3 | 0.0 | 87.4 | 0.0 | 3.1 | 0.9 |
| 45.5 | 0.0 | 116.9 | 0.0 | 0.8 | 0.5 |
| 14.6 | 0.0 | 0.2 | 0.0 | 0.4 | 0.0 |
| 5.9 | 0.0 | 83.5 | 0.0 | | 0.6 |
| 11.1 | | 44.2 | 0.0 | | 0.0 |

Table 2- CD45.2$^+$ B cells/μL in flow spleen sample

| Wt Saline | NSG Saline | Wt PFF | NSG PFF | NSG PFF T | NSG PFF B |
|---|---|---|---|---|---|
| 2855.2 | 0.0 | 0.0 | 0.0 | 0.2 | 3.1 |
| 2061.7 | 0.0 | 3.6 | 0.0 | 2.0 | 20.9 |
| 4495.3 | 0.0 | 0.2 | 0.0 | 4.4 | 1.0 |
| 5662.9 | 0.0 | 0.0 | 0.0 | 0.0 | 20.9 |
| 383.8 | 3.3 | 4571.8 | 0.5 | 0.0 | 39.3 |
| 6711.6 | 0.0 | 2195.2 | 0.7 | 0.0 | 86.8 |
| 5192.7 | 0.0 | 2135.0 | 0.1 | 0.0 | 78.7 |
| 2756.2 | 0.0 | 5168.7 | 0.4 | | 36.3 |
| 3158.1 | | 5605.8 | 0.0 | | 102.5 |
| 2116.1 | | 6673.1 | | | |
| | | 5458.3 | | | |

Table 3- CD45.2$^+$ T cells/μL in flow blood sample

| Wt Saline | NSG Saline | Wt PFF | NSG PFF | NSG PFF T | NSG PFF B |
|---|---|---|---|---|---|
| 7.0 | 0.0 | 0.8 | 0.0 | 81.4 | 0.0 |
| 53.3 | 0.0 | 0.0 | 0.0 | 58.5 | 0.0 |
| 79.3 | 0.0 | 45.3 | 0.0 | 118.1 | 0.0 |
| 16.4 | 0.0 | 0.0 | 0.0 | 111.8 | 0.0 |
| 23.4 | 0.0 | 30.6 | 0.0 | 129.1 | 0.0 |
| 3.9 | 0.0 | 51.0 | 0.0 | 264.9 | 0.0 |
| 1.8 | 0.0 | 0.0 | 0.0 | 391.7 | 0.1 |
| 7.5 | 0.0 | 62.8 | 0.0 | | 0.0 |
| 0.6 | | 22.1 | 0.0 | | 2.0 |
| | | 7.4 | | | |
| | | 16.1 | | | |

Table 4- CD45.2$^+$ T cells/μL in flow spleen sample

| Wt Saline | NSG Saline | Wt PFF | NSG PFF | NSG PFF T | NSG PFF B |
|---|---|---|---|---|---|
| 1174.8 | 0.0 | 17.6 | 0.0 | 432.6 | 0.4 |
| 1103.8 | 0.0 | 0.0 | 0.0 | 398.8 | 15.8 |
| 1843.4 | 0.0 | 479.4 | 0.0 | 79.1 | 78.5 |
| 2508.5 | 0.0 | 0.0 | 0.0 | 100.6 | 0.1 |
| 57.5 | 0.3 | 2194.9 | 0.3 | 101.8 | 0.2 |
| 3631.6 | 0.0 | 1073.2 | 0.5 | 1.8 | 2.1 |
| 3151.6 | 0.0 | 1157.3 | 0.0 | 7.2 | 0.3 |
| 2103.9 | 0.0 | 2244.7 | 0.2 | | 0.0 |
| 1813.6 | | 2409.9 | 0.0 | | 6.0 |
| 1705.6 | | 3235.1 | | | |
| | | 3315.3 | | | |

**Supplementary Figure 1**

### Example Gating Strategy using WT PBS-Treated Mouse

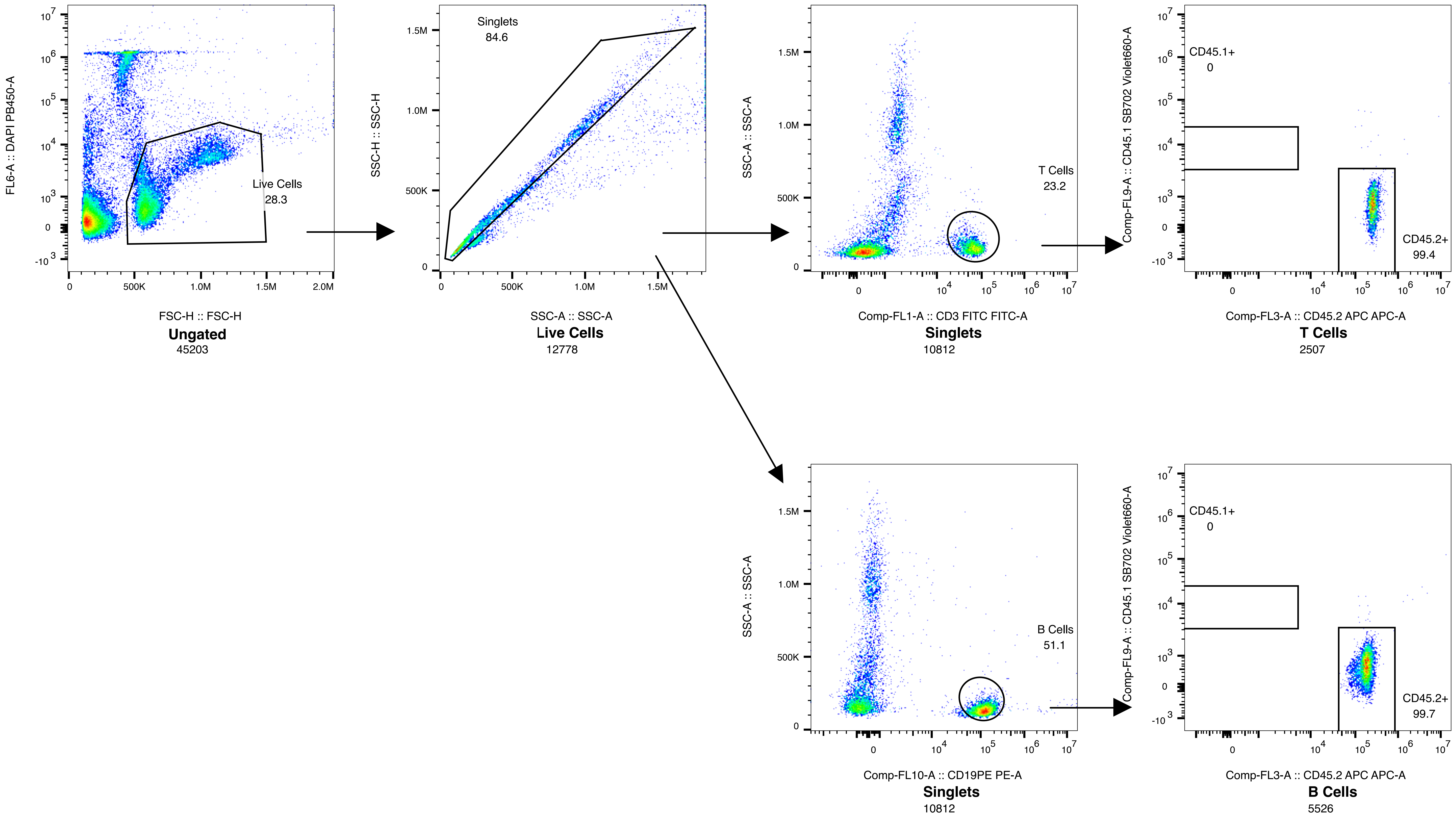

Supplementary 2

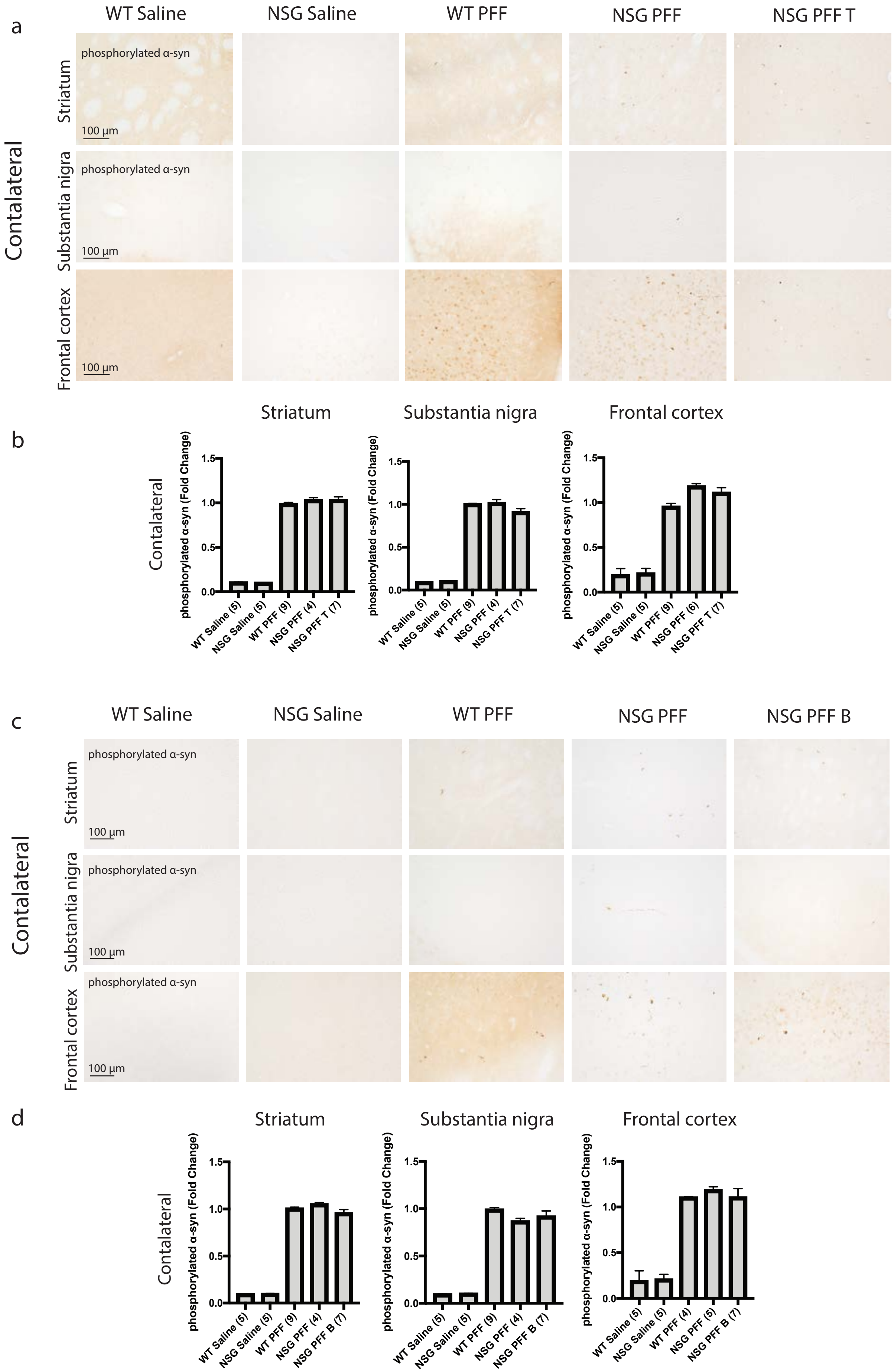

Supplementary Figure 3

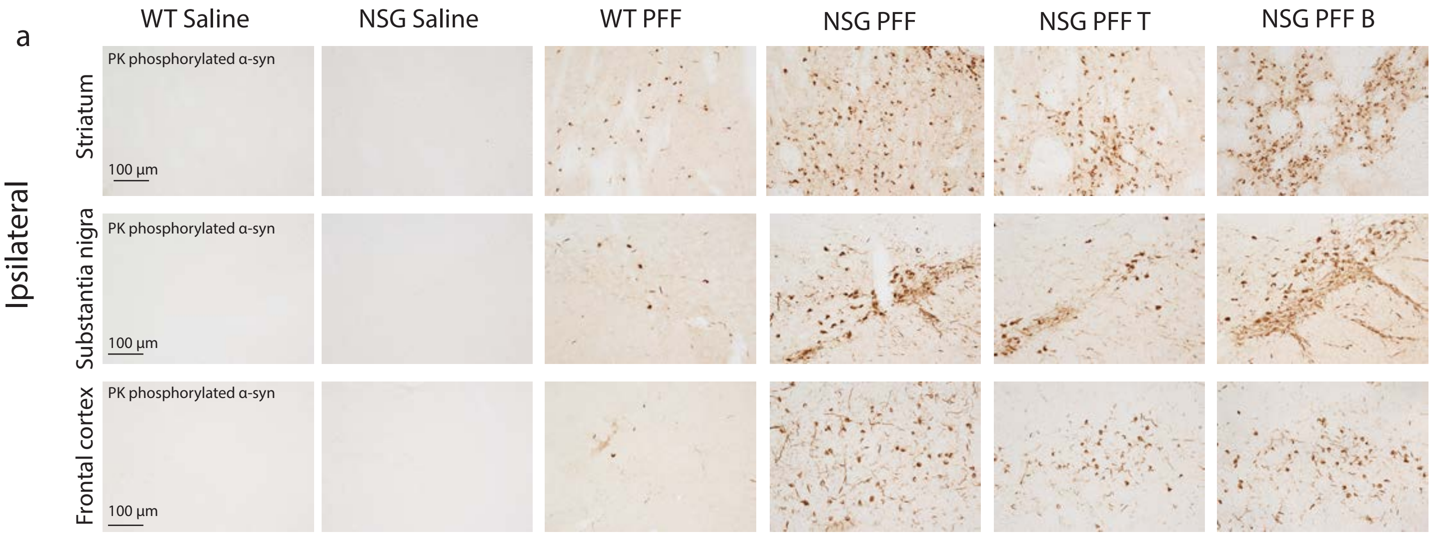
